# Accelerated Simulation of Multi-Electrode Arrays Using Sparse and Low-Rank Matrix Techniques

**DOI:** 10.1101/2024.07.29.605687

**Authors:** Nathan Jensen, Zhijie Charles Chen, Anna Kochnev Goldstein, Daniel Palanker

**Author notes:** The first two authors contributed equally to this work.

## Abstract

**Objective:** Modeling of Multi-Electrode Arrays used in neural stimulation can be computationally challenging since it may involve incredibly dense circuits with millions of interconnected resistors, representing current pathways in an electrolyte (resistance matrix), coupled to nonlinear circuits of the stimulating pixels themselves. Here, we present a method for accelerating the modeling of such circuits with minimal error by using a sparse plus low-rank approximation of the resistance matrix.

**Methods:** We prove that thresholding of the resistance matrix elements enables its sparsification with minimized error. This is accomplished with a sorting algorithm, implying efficient O (N log (N)) complexity. The eigenvalue-based low-rank compensation then helps achieve greater accuracy without significantly increasing the problem size. *Results:* Utilizing these matrix techniques, we reduced the computation time of the simulation of multi-electrode arrays by about 10-fold, while maintaining an average error of less than 0.3% in the current injected from each electrode. We also show a case where acceleration reaches at least 133 times with additional error in the range of 4%, demonstrating the ability of this algorithm to perform under extreme conditions.

**Conclusion:** Critical improvements in the efficiency of simulations of the electric field generated by multi-electrode arrays presented here enable the computational modeling of high-fidelity neural implants with thousands of pixels, previously impossible. *Significance:* Computational acceleration techniques described in this manuscript were developed for simulation of high-resolution photovoltaic retinal prostheses, but they are also immediately applicable to any circuits involving dense connections between nodes, and, with modifications, more generally to any systems involving non-sparse matrices.

## I. Introduction

Retinal prostheses show promise in restoring sight to patients impaired by retinal degeneration, such as age-related macular degeneration (AMD) and retinitis pigmentosa. One such device that has shown particular success in clinical trials is the photovoltaic subretinal system (PRIMA) developed by Science Corp (formerly Pixium Vision). The PRIMA implant uses a 2-D array of photodiodes to convert images projected with near-infrared light into electrical current. When injected into the tissue via a capacitive multi-electrode array, this current generates electric potential, which stimulates retinal bipolar cells. By doing so, the PRIMA implant imitates the neural signals provided by photoreceptors in a healthy retina. This approach restores form vision in patients with retinal degenerative diseases, such as age-related macular degeneration [1].

The current PRIMA implant is 2 mm × 2 mm in size, corresponding to an approximately 7° field of view [2], filled with a hexagonal array of 100 µm wide pixels. The 100 µm gap between lines of a letter, such as a Tumbling E or Landolt C (LDC), corresponds to a visual acuity of 20/420 [1].

It has been demonstrated that grating acuity in rodents matches the pixel size of subretinal photovoltaic implants, up to their natural resolution limit of 28 µm [3]–[5]. It would follow that to increase the resolution in patients beyond 20/420, the size of individual pixels should be decreased (ideally up to 5 µm for 20/20 vision); and to widen the field of view, the implant width should be increased up to some anatomical constraints – such as a size of the geographic atrophy in AMD patients, retinotomy length, etc. Predicting the electric field in electrolyte generated by smaller pixels in a larger array requires an accurate modeling framework for such an implant. For this purpose, the Retinal Prosthesis Simulator (RPSim) has been developed [6].

RPSim is a platform for calculating the electric field in tissue (an electrolytic medium) generated by a multi-electrode array, each pixel of which may include an electric circuit [6]. It does so by coupling the power of finite element method (FEM) physics modelling with the efficiency of SPICE-based circuit solvers. For these respective tasks, RPSim relies on COMSOL Multiphysics® [7] and XYCE, a SPICE-like circuit solver developed by Sandia National Laboratories [8]. First, the elementary field of an electrode and resistance of the medium between the electrodes are calculated with FEM, and then the results are used to build an accurate circuit model, which includes all pixels coupled to a common electrolyte via their electrodes. The circuit model provides the dynamics of an electric current in each pixel, which is then used to calculate the resulting electric field in the medium as a sum of the elementary fields from each electrode, weighted by its respective current.

With the modeling performed in RPSim, the number of computations increases quadratically with the number of pixels on such a photovoltaic array due to a cross-coupling between each electrode in the model. The number of electrodes, *N*, that can be tiled on a circular implant depends on the width of individual pixels, *W*, and the diameter of the implant *D*. For example, more than 20,000 pixels of 20 µm fit on a 3 mm wide implant. The number of resistors in a mesh representing the electrolyte exceeds 4 × 10^8^. As pixels get smaller, and the implant – larger, computations quickly become unattainably long, and require an incredibly large operating memory, which prevents efficient modelling of the next-generation implants. Dramatically simplifying the simulation of subretinal implants is necessary to enable the simulation of implants with tens of thousands of electrodes, which requires too much memory even for modern desktops. Additionally, it can dramatically decrease the simulation time, which enables iterative optimization of the design of the next generation devices.

As mentioned, to solve the circuit dynamics involved in a neurostimulating array, RPSim relies on the circuit solver Xyce [6], [8]. One would be hard pressed to develop a tool that can outperform such a mature and proven simulator, capable of handling millions of circuit elements. However, these solvers have been developed using a node-based approach and optimized under the assumption that the matrix between adjacent nodes is sparse. This assumption breaks down for mutually coupled multi-electrode arrays that suffer from an extremely dense conductance matrix. To help, one can find a sparse matrix approximation, but the error introduced by such a technique can be very large. We can, however, reduce this error by using low rank compensation matrices. To the author’s knowledge, circuit solvers, such as Xyce, have no such built-in methods to increase their computational efficiency. Instead, for such large-scale problems they suggest the user rely on parallel computing and iterative methods, such as Krylov, which require a well-designed and problem-specific preconditioner for good performance [9]–[11]. Here, we present a much simpler and more astute alternative to such complicated algorithms for reducing the computational intensity of modeling a large mesh network of photovoltaic pixels connected via a common electrolyte. This approach is generalizable to other massive electronic circuits coupled to an electrolyte or another conductive medium and is specifically useful for high precision modeling of multi-electrode arrays with thousands of pixels. For example, Neuralink’s volumetric brain-machine interface contains 3072 electrodes [12]. Precision Neuroscience has also developed a cortical surface interface with thousands of electrodes [13].

Iterative methods, including Krylov based ones, work well when the matrices are sparse. They also require large amounts of memory and can outright fail due to not converging or if the problem contains too large of an irreducible block. Performance of such methods depends largely on the combination of the circuit, the solver, and the preconditioner. Failure to set these correctly will harm performance at best or prevent convergence at worst. When successful, iterative methods report simulation speed-ups ranging from 3 to 21 times [14]. Because our method focuses on simplifying the problem rather than improving the solver, it can also be used in conjunction with iterative techniques. In other words, with the method presented here, one can first find a suitable sparse matrix approximation of the circuit and then use that form with iterative techniques, such as Krylov-based solvers, which inherently require sparse matrices for good performance.

## II. Methods

### A. Sparse Approximation

The photovoltaic multielectrode array used in a subretinal implant, such as PRIMA, consists of tiled hexagonal pixels, each of which has one or more photodiodes connected between the active and return electrodes (Fig. 1. Active electrodes are at the center of the pixels, while return electrodes cover the edges of each pixel, connected together over the whole mesh [6]. These photodiodes convert near-infrared light projected from the augmented reality glasses into electric current, which is injected into the retina to stimulate the nearby neurons [2]. All pixels are coupled to each other via a conductive (electrolytic) medium, as shown in Fig. 1, and thus each affects the electrical dynamics of all others. Consider the effects of the pixel 2 – which we will briefly assume has a hemispherical electrode injecting a current *I*_2_ while all other pixels remain idle. A hemispherical electric field will form in the medium, resulting in an elevated electric potential: Δ*V ≈* (*I*_2 ·_ *ρ*)*/*(2*πl*), where *l* is the distance from the center of that electrode and *ρ* is resistivity of the medium. Using this, we can characterize the conductive medium as an ohmic resistance between the active pixel 2 and another pixel *i*, which is inversely proportional to the distance between their centers *l*_*i*,2_

**Fig. 1:**
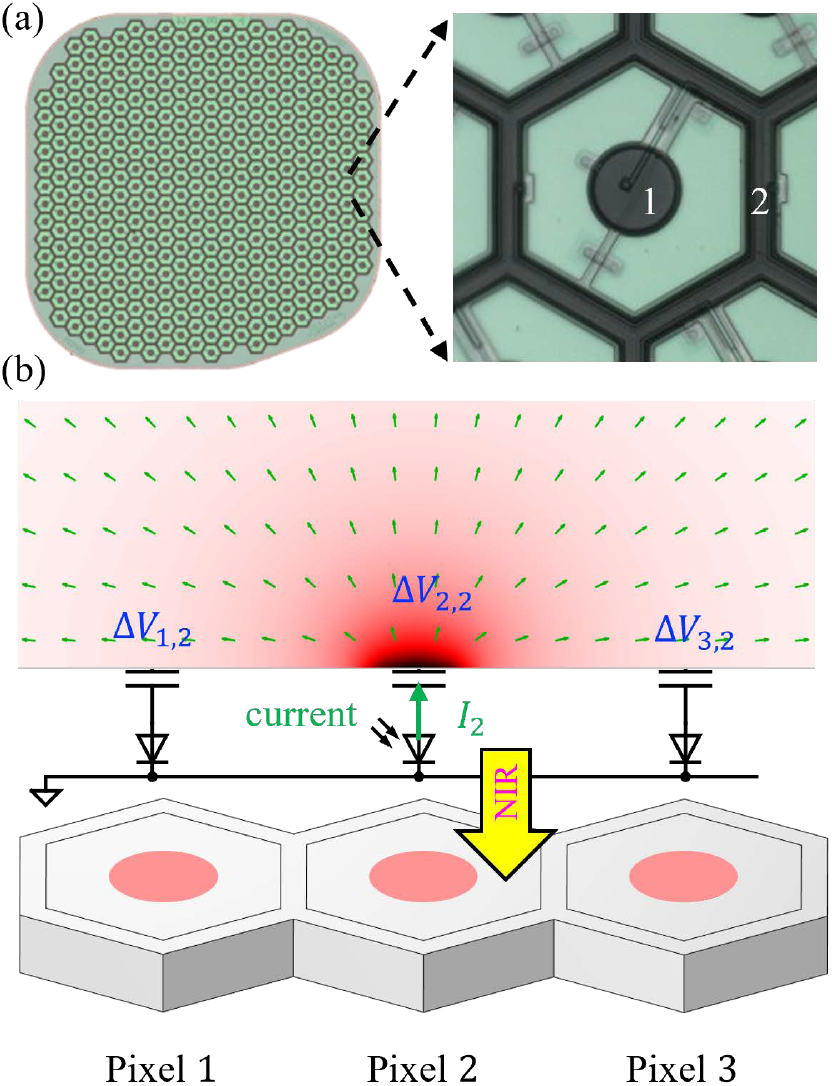
(a) PRIMA implant and magnified view of a single pixel: 1 is the active electrode and 2 is the return electrode. In between the two is the photosensitive area. (b) Visualization of the electric potential and current in electrolyte. The center pixel is illuminated by near-infrared (NIR) light and injects a current *I*_2_, which flows opposite to the photodiode direction.

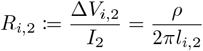

Or more generally,

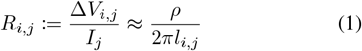

For an adjacent pixel 1, *l*_1,2_ would be on the order of the pixel width *W*. Additionally, for the “self-resistance” of a pixel *R*_*j,j*_ = Δ*V*_*j,j*_*/I*_*j*_ *≈ρ/*(4*r*), where *r* is the electrode radius– following the conventional definition for access resistance of the disk electrode [15]. Furthermore, due to linearity of the conductive medium, when multiple pixels inject current simultaneously, the potential rise Δ*V* at pixel *j* is simply the sum of all the potentials calculated individually, i.e. superposition. This formulation allows modelling the cross-coupling of all *N* electrodes as a mesh network of resistors in matrix form [3], [6].

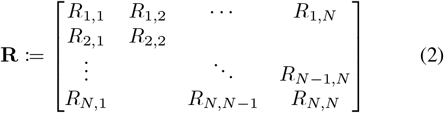

By reciprocity of electromagnetism, this matrix will be symmetric. The size of this resistor mesh scales quadratically with the number of pixels, and hence it scales to the fourth power with both the implant width as well as the inverse of the pixel size. Consequently, the computational workload of simulating such a mesh grows accordingly for both factors. This mesh is the primary contributing factor to the polynomial relationship between computational complexity and the number of pixels on an implant.

The problem of reducing the computational complexity can be framed as finding a matrix which approximates the conductance matrix **G** := **R**^*−*1^, but is in some way computationally lighter. For example, one may find a sparse approximation, denoted as **S**. Because of the spatial dependence of *R*_*i,j*_ – being inversely proportional to *l*_*i,j*_ – one intuitive method of sparsification would be to find some threshold value for the distance between two electrodes, such that their coupling is so small it can be neglected, i.e. set to 0 in the conductance matrix. However, the total number of coupled electrodes, *η*, some distance *l*_*i,j*_ away from the center electrode is roughly proportional to *l*_*i,j*_

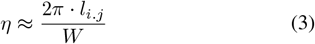

where *W* is the width of a pixel.

The variable *l*_*i,j*_ in (2) and (3) cancels out, such that the overall contribution to the effective access resistance of one electrode from consecutive rings of neighboring electrodes is largely constant with distance. Therefore, this naïve thresholding alone can result in a very large error, which will be addressed later. Formally, one needs to find a sparse approximation **S** for the network **G** with no more than *k* non-zero entries, such that **S** is positive definite and the error

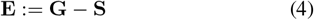

is minimized, where *k* is a value dependent on the computational resources available. To minimize the error, which is expressed in matrix form here, we will minimize the spectral energy— defined as the sum of squares of all the eigenvalues (*λ*) of **E**, which for a symmetric matrix is equivalent to the square of the Frobenius norm of **E**.

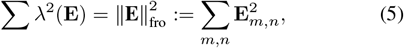

where ()_*m,n*_ denotes the entry in the *m*th row and the *n*th column of a matrix. Therefore, combining (4) and (5), this can be written as the optimization problem:

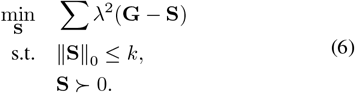

This problem has a direct solution, which does not require iteration to solve. Let *I*^+^ be a set comprising all indices (*m, n*) such that **S**_*m,n*_≠0, and having complementary set *I*^*−*^ := {(*m, n*)|**S**_*m,n*_ = 0}. Note that

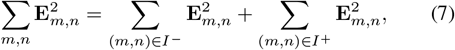

and for (*m, n*) ∈ *I*^*−*^, **E**_(*m,n*)_ = **G**_(*m,n*)_. From this One has:

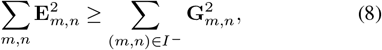

where the equality is achieved if and only if **S**_*m,n*_ = **G**_*m,n*_ for all (*m, n*) ∈ *I*^+^. In this case,

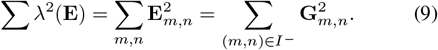

The solution to this optimization is indeed found by thresh-olding **G** by it’s *k*^*th*^ largest entries, as postulated above, and setting all other values in **G** to be 0. Thresholding can be accomplished using a sorting algorithm, such as the popular quicksort, which has an average complexity time in Big O notation *O*(*N* log (*N*)) [16].

If we define a column vector of the potential at each pixel as ***V***, then from multidimensional Ohm’s Law, we can use the error in the conductance matrix, **E**, to define the vector containing the error in injected current for each pixel as ***I***_err_ := **E*V***. This error is therefore bounded by:

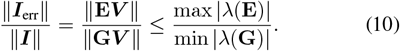

### B. Low Rank Compensation

By this metric, the maximum eigenvalue in **E** contributes the most to the current error, and because of this, error might be decreased by compensating for the specific terms in **E**. The largest eigenvalue of **E** represents the principal component of the error – and represents a relatively uniform current profile caused by the error of neglecting small-magnitude couplings between distant electrodes, as was discussed. This error is approximately the same for each electrode in the array, albeit slightly different for electrodes closer to the edge of the implant that have fewer close neighbors. To correct this, a simple error-reduction technique will be introduced and deemed general compensation since the error from these principal components does not involve any information about which electrodes are injecting current. This compensation is achieved through the addition of a low rank matrix to the sparse matrix **S** and having error **E**_1_:

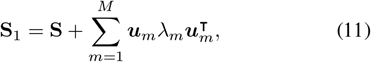

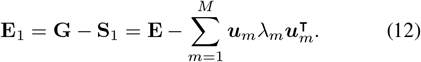

where **S**_1_ is the resulting compensated matrix, *λ*_*m*_ is the *m*th largest eigenvalue of **E**, and ***u***_*m*_ is the corresponding eigenvector. *M* is thus the total number of desired compensation vectors, which can be as large as the number of eigenvalues in the error matrix **E**. By definition, this will be at most the number of electrodes *N*. As *M* increases, the total error will decrease at the cost of a larger compensation matrix. The resulting compensation mode from the principal component is largely uniform as thresholding removes a similar number of interconnects for each electrode, removal of which results in the equivalent resistance of an electrode being too high. To rectify that, the compensation must provide additional conductivity to the electrode. On the edges of the array, less interconnects are removed from the matrix and thus less added conductivity is needed. This gives rise to the pattern shown in Fig. 2(a).

**Fig. 2:**
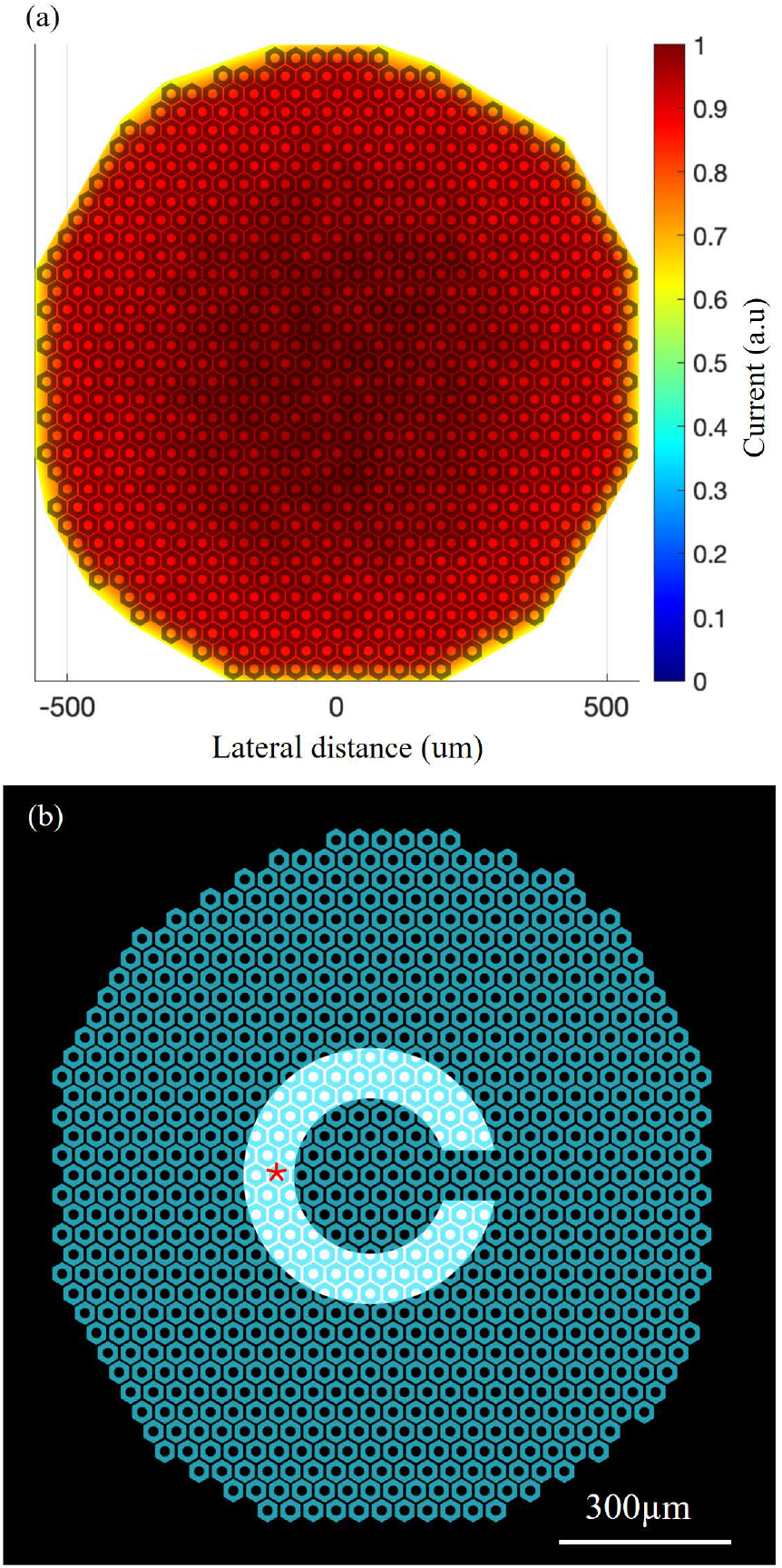
(a) Map of the principal component of the error matrix **E** resulting from thresholding out 90% of the smallest magnitude entries of **G**, the corresponding conductance matrix for a 40 µm monopolar implant. Values are normalized by the largest entry. (b) A Landolt C pattern with a 90 µm wide gap projected onto the array. The red asterisk labels the pixel whose current is plotted in Fig. 5(a).

Further specific compensation terms can be added with knowledge of the illumination patterns of interest or the operation mode of the implant, such as if there is no forward conduction through the diodes, which is called a current limited regime [14]. For example, if a known pattern is projected, resulting in an expected photocurrent in each pixel ***I***_*pho*_, then the expected voltage distribution can be calculated as ***v*** = **G**^*−*1^***I***_pho_, assuming that the device operates under current limited conditions. The corresponding current error is ***w*** = **E**_1_***v***. Since ***v*** constitutes the primary voltage mode from the pattern, ***w*** will be the primary current error. Therefore, we can add the following specific compensation:

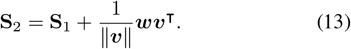

This image-specific compensation is generalizable such that successive low-rank compensations can be added for a variety of illumination patterns and implant operation conditions.

### C. Circuit Implementation

In RPSim, the conductance matrix is implemented as a mesh network of resistors. With a sparse matrix, we can eliminate some resistors from the model, and as such, implementation involves simply replacing **G** with **S**. However, it would be counterproductive to replace **G** with **S**_1_ or **S**_2_ as the compensation terms are not sparse. They are low rank, but SPICE based circuit solvers do not have support for low-rank representation of matrices. Thus, maintaining computational efficiency while introducing compensation terms requires a means of adding a low-rank matrix to a sparse one without sacrificing the sparsity. This can be accomplished in Xyce through the addition of several additional nodes.

Let the rank-1 compensation matrix be ***w****λ****v***^⊺^. The implant has *N* electrodes, including the active electrodes and the return units. ***w*** = [*w*_1_, *w*_2_, · · ·, *w*_*N*_]^⊺^ and ***v*** = [*v*_1_, *v*_2_, · · ·, *v*_*N*_]^⊺^ are *N* -dimensional vectors. For the general compensation, ***w*** = ***v***; for the specific compensation, *λ* = 1*/*∥ ***v*∥**. One may implement the compensation ***w****λ****v***^⊺^ with 2*N* voltage-controlled current sources (VCCSs) and a resistor, as illustrated in Fig. 3. Each VCCS in the left group takes voltage control from an electrode, multiplies the voltage with the corresponding entry in ***v***, and adds the current to a resistor *λ*. The voltage at the top node of the resistor is *λ****v***^⊺^***V***. The VCCSs in the right group then take the voltage from the top node of *λ*, multiply it with ***w***, and pull the corresponding currents from the same set of electrodes 1 to *N*. The coupled *i − v* relationship between all electrodes is given by

**Fig. 3:**
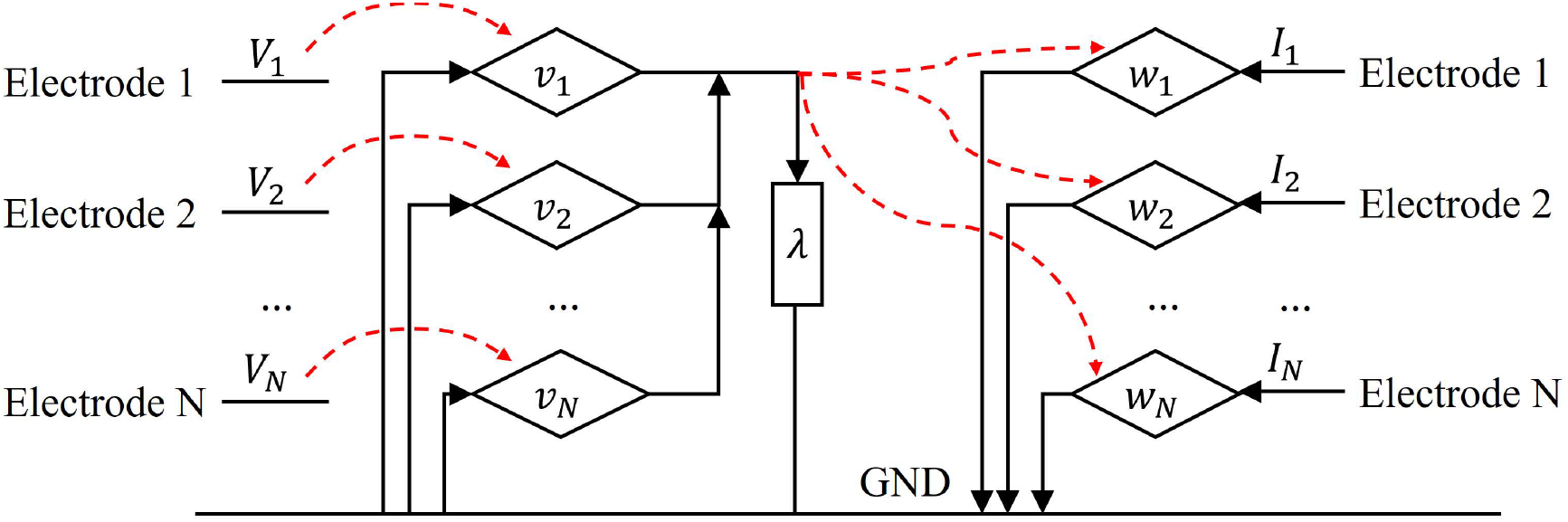
Circuit level implementation of a rank-1 compensation to a sparse conductance matrix. The red dash arrows indicate where each voltage-controlled current source (diamond) takes the control signal from, and the rectangle labeled *λ* is a resistor.

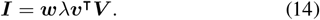

This approach can implement a rank-1 compensation matrix ***w****λ****v***^⊺^ in Xyce, using only 2*N* + 1 elements and one additional node, instead of the (*N* (*N −* 1))*/*2 resistors otherwise required by the full resistor mesh. For a more formal proof of this concept, consider a *N × N* sparse matrix **S** and a rank-*L* compensation term **WΛV**^⊺^, where **W** and **V** have dimensions of *N* × *L*, **Λ** is a *L* × *L* diagonal matrix, and *L* ≪ *N*. Rather than explicitly solving a linear system with an unknown vector variable ***x*** and a known vector ***b***:

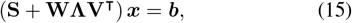

which enjoys neither sparsity nor low-rankness, one can instead solve the following system with *L* additional dimensions:

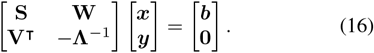

The augmented equation above requires

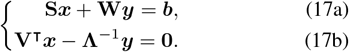

Equation (17b) gives:

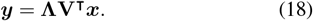

Substituting the above ***y*** into (17a) results in (15), while the matrix in (16) maintains its sparsity, demonstrating the validity of such an approach.

### D. Additional Considerations

Further simplifications of the circuit can be made under certain conditions. For example, under low irradiances when the implant operates in the current limited regime, the voltage across the parallel diode used in a conventional model of a photodiode will be below the threshold of forward diode conduction. In this case, the forward current through the diode will be negligible and we can model the diode as an open circuit without sacrificing accuracy. This is advantageous as diodes are the only non-linear component in the circuit and removing them can significantly increase the computation speed. This technique can be especially useful for modelling larger photovoltaic arrays with smaller pixels at low irradiances. However, this assumption will quickly break down at high irradiances when the diodes begin to conduct. In the context of subretinal implants, the maximum irradiance level before this occurs also depends on the number of diodes in series within a pixel, typically ranging from 1 to 3 [17].

## III. Results and Discussion

For demonstration purposes, a 1.5 mm implant composed of *N* = 821 hexagonal 40 µm wide monopolar pixels was modelled. For sparsification, the thresholding value *k* was set in a ratiometric manner with the ratio K, such that *k* = ∥**G**∥ _0_ · K. Here, K was set to 0.1, which corresponds to keeping 10% of the initial entries of **G** – or a 90% reduction in size of the resistor mesh. Recall that error is bounded by the maximum eigenvalue of **E** divided by the minimum eigenvalue of **G** (10), plotted in Fig. 4(a). The initial bound on the error – (10) evaluated at the first eigenvalue in Fig. 4(a) – is about 70%. However, this error can be dramatically improved by compensating for just the first principal component of the error. In this case, after implementing the general compensation with the eigenvector associated with the largest eigenvalue of **E** (*M* = 1), the error will be bounded by the second largest eigenvalue in **E**, with an error of less than 7%. This demonstrates that we can improve the bounded error by an order of magnitude by adding a rank one compensation matrix, which corresponds to 2*N* +1 circuit elements. Doing so dramatically reduced the super-linear scaling in computational time, such that one can choose an acceptable error and runtime. With these parameters, the original netlist size was 13 MB, while after simplification and compensation it decreased to 1.7 MB.

**Fig. 4:**
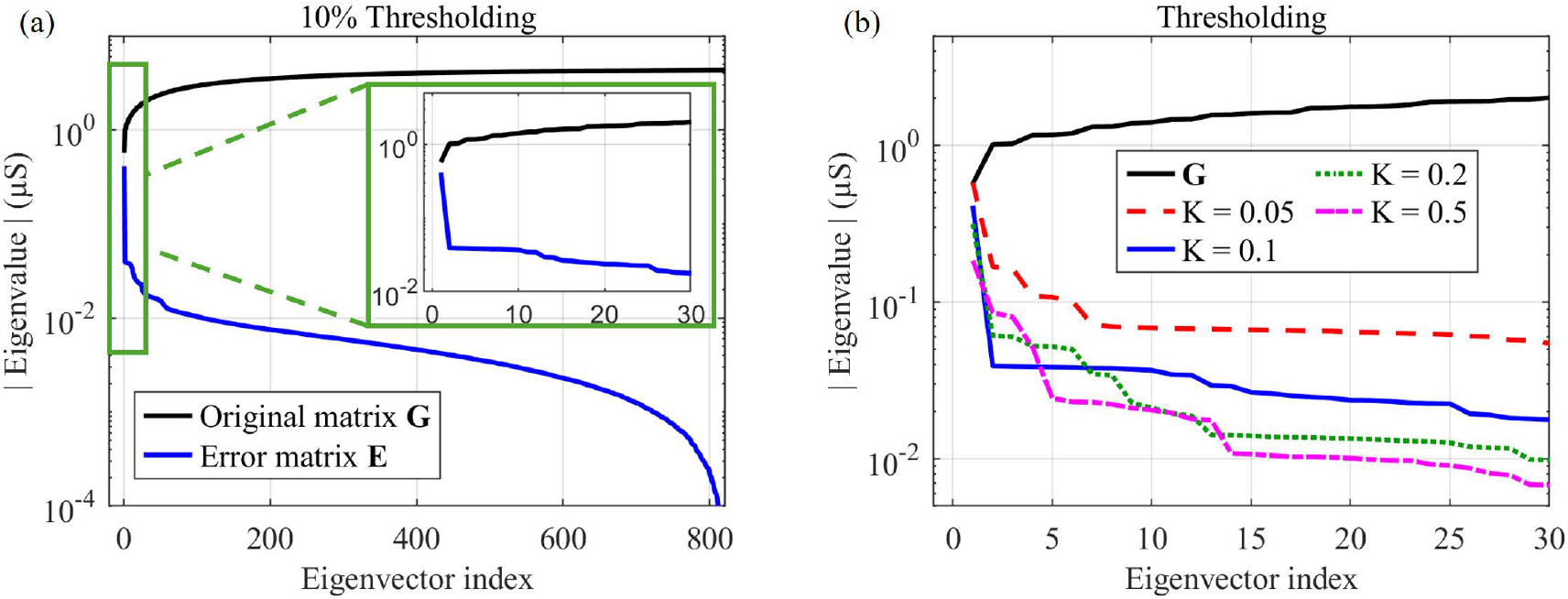
Eigenvalues of the conductance matrix **G** sorted by ascending magnitudes, and those of the error matrix **E** by descending magnitudes, **E** = **G − S**. The relative error is bounded by the inverse of the gap between the two spectra. (a) **S** is a sparse approximation of **G** with K = 0.1, the parameter used for simulations in this paper. (b) The error spectrum is shown for several values of K.

It is important to note that the error spectrum in Fig. 4 depends on the implant geometry as well as the value of K. In some cases, more than the first principal component must be included to achieve an acceptable error, at the cost of adding a corresponding number of circuit elements to the simulation, as illustrated in Fig. 4(b). Notably, for certain values of *M*, choosing a lower value of K could simultaneously decrease both the error and the simulation time. This shows the importance of monitoring the spectrum of **G** and **E** when selecting these parameters, as otherwise one could select a suboptimal combination. For larger values of *M*, i.e. *>* 13 for the ratios plotted in Fig. 4(b), this is no longer true and a smaller value of K will always result in a larger error.

To calculate the actual error in electric current resulting from the thresholding and compensation described above, a Landolt C pattern with a 90 µm wide gap shown in Fig. 2(b) was projected at an irradiance level of 3 mW*/*mm^2^, with a pulse width of 9.8 ms, repeated at 30 Hz. The simulation was 330 ms long, enough time to allow convergence to steady state. The simulation was ran using the unmodified conductance matrix **G**, an uncompensated sparse matrix **S**, the sparse matrix with general compensation **S**_1_ and the sparse matrix with both general and image-specific compensation terms **S**_2_. For each case, the current injection from each pixel is found using RPSim, and then, with the unmodified (full) matrix serving as a benchmark, the relative error was calculated for both the 2-norm of currents from all pixels and the maximum current from all pixels. This error is calculated from the respective currents (2-norm or maximum) when using the original full matrix, *i*_*full*_, and the simplified matrix *i*_*simp*_. Relative error is thus found as (*i*_*full −*_ *i*_*simp*_)*/i*_*simp*_. The results are shown in Fig. 5. Over the plotted window of the simulation, the average of the 2-norm of the relative error for the uncompensated sparse matrix is about 4.6%, while with the general compensation term this is reduced to 0.65%, and with the image-specific compensation - to 0.036%. The results of timing and the normalized root mean square (NRMS) error

**Fig. 5:**
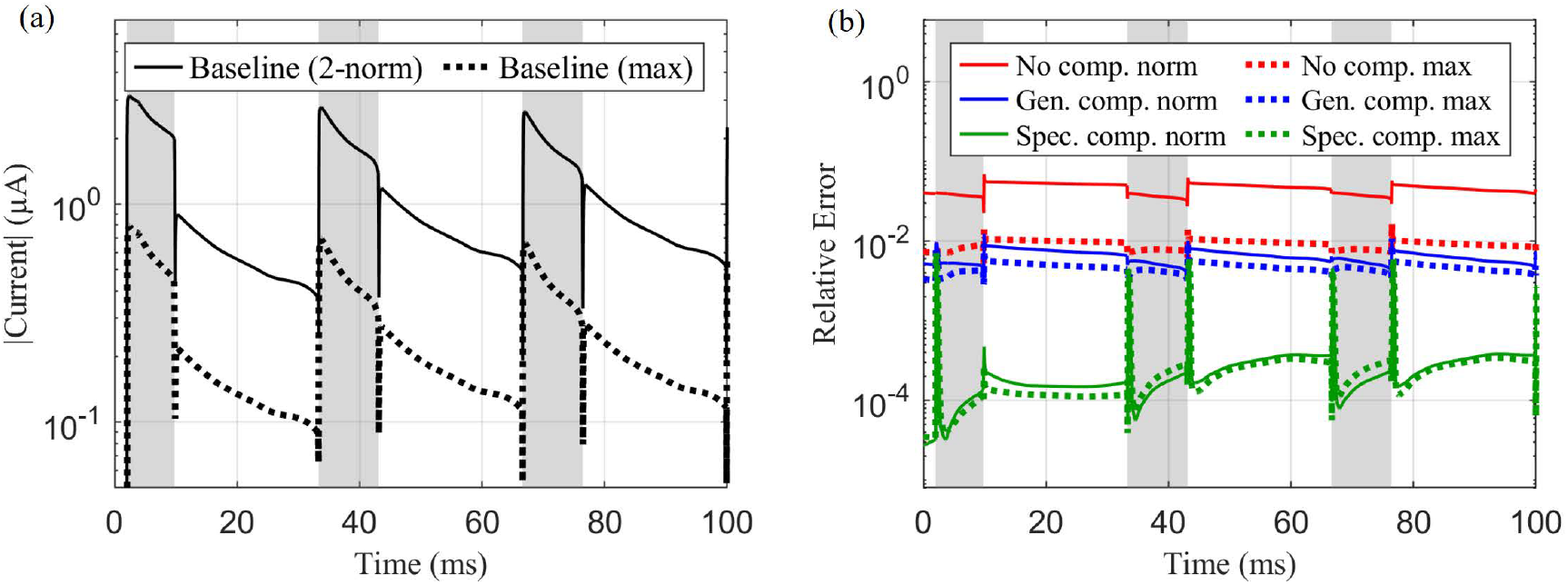
Currents on the implant across time for the stimulation shown in Fig. 2, the shaded portion indicates the times when the stimulation pattern is turned on. (a) Baseline calculated with unmodified full matrix. Maximum current injected from the pixel shown in Fig. 2(b) (max) and the 2-norm of the currents of all pixels. (b) Relative error of the pixel shown in Fig. 2(b) (max) and the 2-norm of the error of the currents for all pixels: for the uncompensated matrix, with general compensation, and with image-specific compensation.

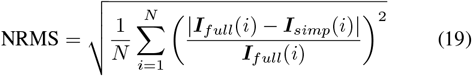

of the average current during an illumination pulse are recorded in Table I. With only the primary component (*M* = 1), compensation adds a negligible amount of computation time but reduces the error more than 10 times. Similarly, the image-specific compensation adds very little computation time while providing an equally large reduction in error. Illustrations of the error distribution for these different simulations are provided in Fig. 7

**TABLE 1.** The simulation time and error in the average current for all pixels during an illumination pulse (Eqaution 19) for a Landolt C pattern (Fig. 2(a)) projected at 30 Hz during 330 ms. **S** was calculated using K = 0.1, **S**_1_ adds compensation with *M* = 1, and **S**_2_ further adds image-specific compensation.

| Conductance Matrix | Simulation Time (minutes) | NRMS Error |
| --- | --- | --- |
| G | 5.44 | N/A |
| S | 0.63 | 71.6% |
| S1 | 0.64 | 3.16% |
| S2 | 0.68 | 0.18% |

Additionally, to confirm the tractability of this method, the simulation was run with *M* = 1 and K = 0.1 for various implant geometries – bipolar and monopolar devices including 100, 75, 40, and 20 µm pixel sizes. The number of pixels on these devices ranges from 205 to 2806. Bipolar devices have both an active and return electrode on every pixel, whereas monopolar devices have one shared return electrode along the edge of the device [6]. This means that bipolar devices have nearly twice as many electrodes, and therefore a 4x larger resistance mesh compared to a monopolar implant of the same dimensions. We can see in Fig. 6(a) that in all cases the computation time was reduced by the expected factor of 10, corresponding to the thresholding ratio. To enable simulation within a reasonable timeframe for larger implants with smaller pixels, the thresholding value can be reduced even further.

**Fig. 6:**
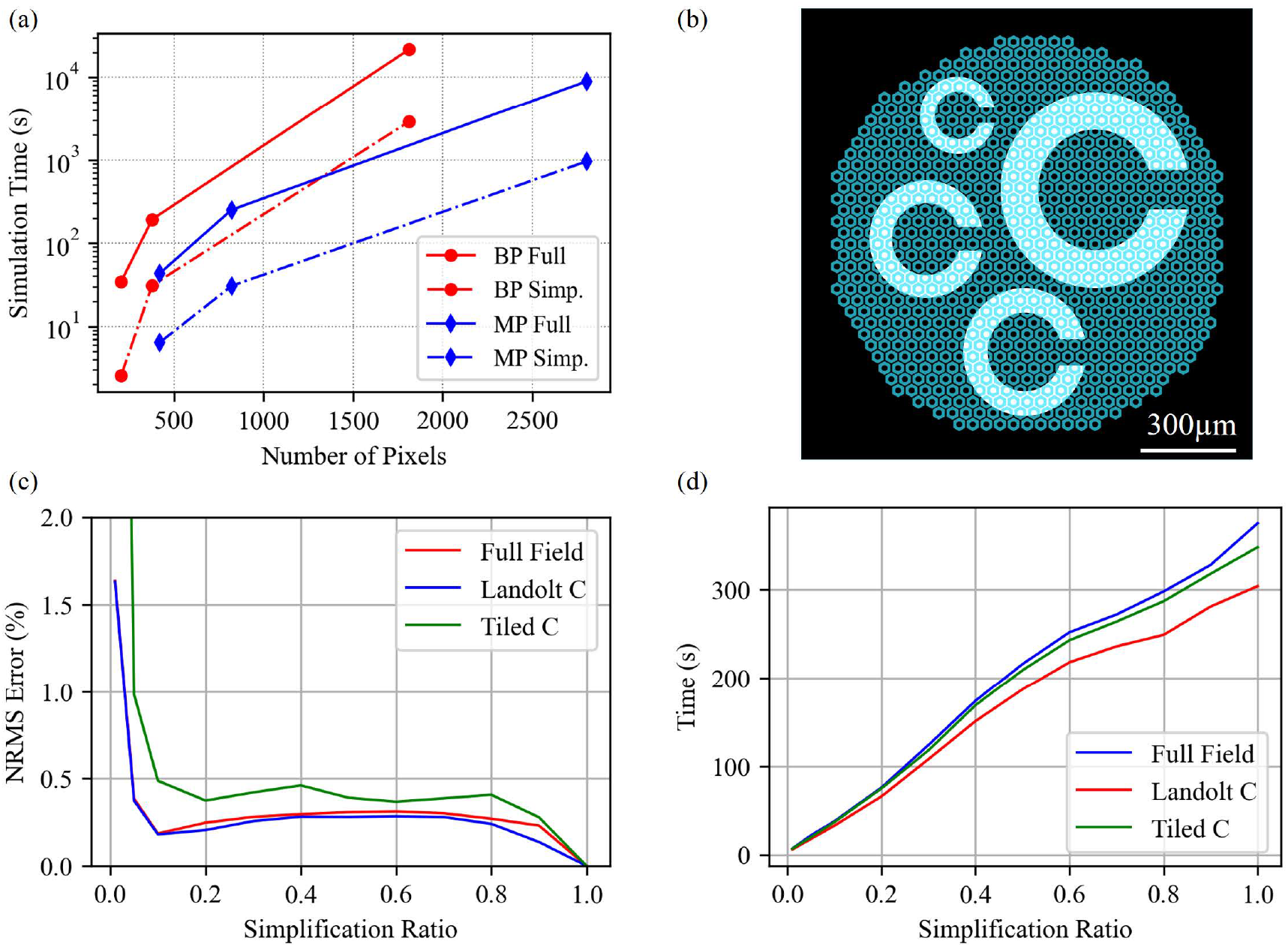
(a) Simulation time for the full conductance matrix and the simplified and compensated matrix for bipolar (BP) and monopolar (MP) devices. (b) ‘Tiled C’ pattern used in (c) and (d), ‘Landolt C’ is shown in Fig. 2(b) and ‘Full Field’ means that all pixels are uniformly illuminated. (c) Error for various patterns as a function of the simplification ratio K, which changes the amount of simplification for the conductance matrix. The compensation was kept at *M* = 1 for all cases. (d) Simulation time as a function of K for various patterns.

**Fig. 7:**
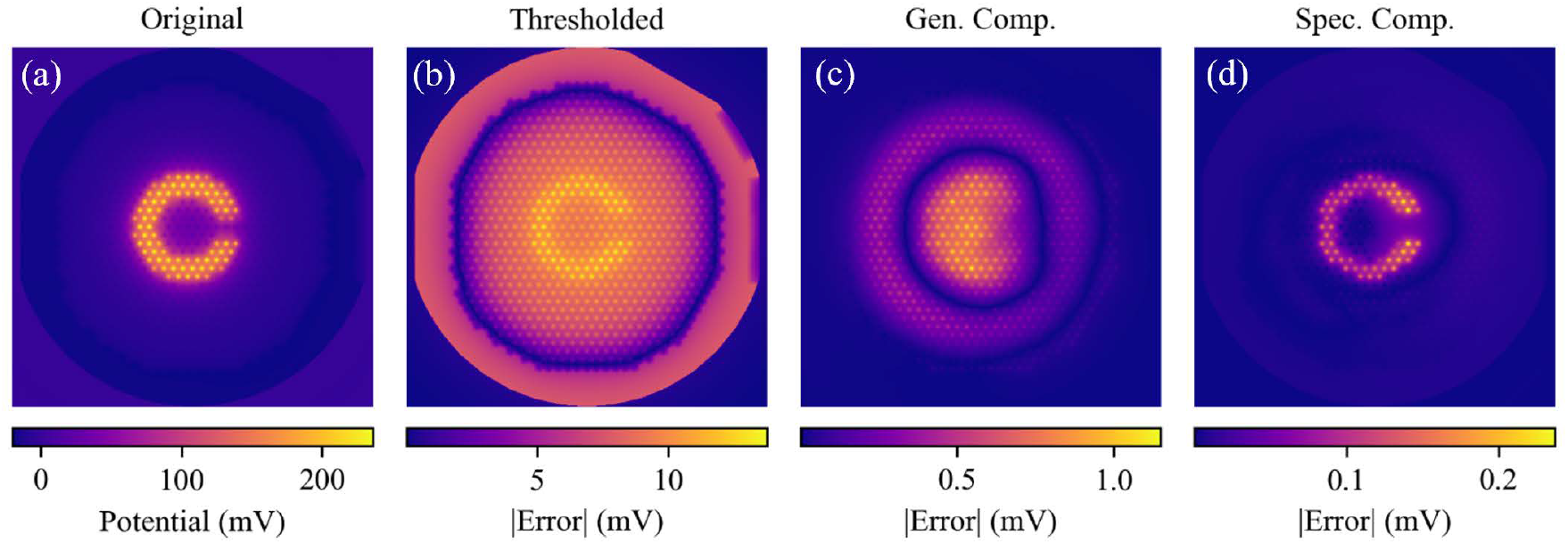
(a) Electric potential at the surface of the implant illuminated with the pattern shown in Fig. 2(b), computed with unmodified matrix **G**. (b-d) Absolute value of the error in electric potential computed at the following settings: (b) thresholded matrix with K = 0.1, (c) with compensation for the first principal component *M* = 1, (d) with additional image-specific compensation.

Such adjustments will become necessary for higher electrode counts as the netlist size scales quadratically with the number of electrodes. The 1.5 mm-wide monopolar implant with 821 pixels of 40 µm totals 3.4 × 10^5^ elements in the resistor mesh, a netlist file of 17 MB, and about 6 min of CPU time. For the same size monopolar implant composed of 2806 pixels of 20 µm, RPSim computes with about 4 × 10^6^ elements, a 220 MB netlist file, and around 3 hours of CPU time. Based on such scaling, a 3 mm× 4 mm large implant would contain about 2 × 10^8^ elements, an 11 GB netlist, and an intractable CPU time on a typical computer, especially as the computational time usually grows super-linearly with the problem scale.

To evaluate performance of the algorithm with respect to the thresholding ratio, two additional patterns – first a full field illumination and second several Landolt Cs tiled on the implant (Fig. 6(b)) - were tested on a monopolar 40 µm array, at the same irradiance, frequency, and pulse width as the original pattern (Landolt C with a 90 µm gap size). This was performed for conductance matrices with simplification ratios in the range 0.01 *<* K *<* 1.0 and compensation of the first principal component (*M* = 1) The NRMS error of the average current during an illumination pulse, as well as the simulation time are plotted for each pattern in Fig. 6(c) and (d), respectively. K = 1.0 corresponds to the original complete conductance matrix, which was used as a reference for the error calculation. For all patterns tested, the error is less than 1% for all K *>* 0.05. Additionally, for a simplification ratio of 0.1, the average error for the three patterns is 0.28%, with a corresponding average reduction of the simulation time of 89.5% (9.5x faster). If a lower error or faster simulation is desired, additional low-rank general compensation vectors of the error matrix should be included. The error associated with a specific thresholding ratio will vary with the number of compensation vectors *M*, as illustrated by Fig. 4(b). Additionally, because each illuminated pixel is a current source in the circuit simulation, larger patterns take longer to simulate – explaining the difference in simulation time between patterns in Fig. 6(d). For all simulations, there are elements of computation that do not involve the resistance mesh. As such, there is a small offset in simulation time such that a sparsification ratio of K = 0 will still have a nonzero simulation time. Nonetheless, it is clear from Fig. 6(d) that the computation time for this implant grows linearly with the simplification ratio K, which confirms the postulate that simulation speed is limited primarily by the size of the conductance matrix.

For moderately sized problems, the algorithm presented here allows a tenfold simulation speedup without sacrificing accuracy. This is fast and accurate enough for iterative optimization of implants with about 1,000 electrodes. However, for arrays containing around 10,000 electrodes, careful selection of the hyperparameters is important for good performance. When using this method for optimization, the spectrum of the error should be monitored to select the fastest possible simulation within the desired error. In the context of retinal implants, maximum error should not exceed 5%. This is because stimulation thresholds of retinal bipolar cells are around 10mV, with the PRIMA implant providing 30-60mV across the cell. An error of a couple of millivolts is within patient-to-patient variability and would not affect the resulting prosthetic vision. Precision in other applications, such as cortical brain-machine interfaces, may be higher or lower, and the hyperparameters must be selected accordingly.

To illustrate the aforementioned ability of this algorithm to handle more extreme conditions, a larger bipolar implant with 20 µm pixels, having 7658 electrodes, was tested with the LDC pattern described earlier. With a netlist that itself is over 2GB involving nearly 59 million resistors, simulation of such a device without the techniques presented here would be impossible without access to extreme computational resources. With a 10x simplification slightly modified from that described above – K = 0.1 and *M* = 34 – the simulation was completed in 28.41 hours. While this capability is encouraging, for iterative optimization of the device design, this is still prohibitively long. For further simplification a more optimal set of hyperparameters was found to be K = 0.025 and *M* = 200. The runtime decreased an additional 34.22 times – down to 0.83 hours. The simulation with the full conductance matrix was not completed, but extrapolating the runtime by the number of electrodes would predict it to be around 111 hours. It is important to note that such extrapolation is inherently unreliable: for example, using the same extrapolation based on the simulations with 10 times simplified matrices predicts a runtime of 9.33 hours – a third of the observed 28.41 hours. Therefore, this is likely an overly conservative estimate, and even with that, this would imply a total speedup of 133 times. Similarly, the error with respect to the unmodified simulation is unknown; however, when compared to the simulation using K = 0.1 and *M* = 34, the norm of the average deviation in electrode current was 3.88%.

Finally, To quantify the irradiances at which the reverse conduction through photodiodes can be neglected, a full field and grating patterns with 1-pixel wide bars were tested at 30 Hz with a pulse width of 9.8 ms. The average error (NRMS) in the calculated current, as previously defined, is shown for both the active and return electrodes as a function of irradiance in Fig. 8. The error is introduced once there is a significant voltage across a diode, such that it begins to conduct current. For PRIMA bipolar pixels, there are two photodiodes in series requiring twice the voltage before forward conduction begins. For this reason, the error at a given irradiance is lower for these pixels than for their monopolar counterparts. Additionally, the nominally small inter-pixel coupling for bipolar pixels results in close matching between the error in an active electrode and its local return electrode. This correlation becomes much more complicated for a monopolar design with just one global return electrode. Therefore, in monopolar devices, while the errors for the active and return electrodes do follow similar trends, they still differ significantly. For such reasons, the maximum irradiance introducing less than 1% error is dependent on the pattern and pixel design, but in the worst case for 100 µm PRIMA bipolar pixels this is 2.5 mW*/*mm^2^, while for a monopolar 40 µm device – it is 1.0 mW*/*mm^2^.

**Fig. 8:**
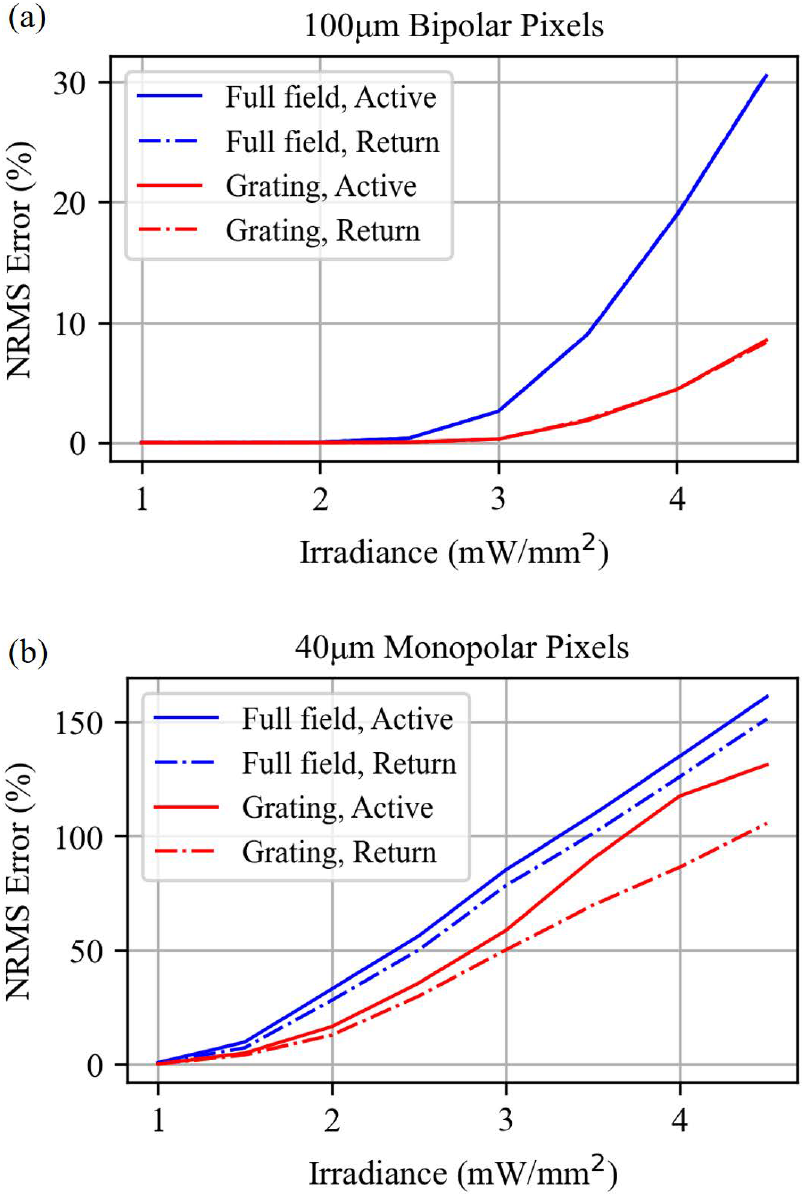
Error introduced in the active and return electrodes by removing diodes from the circuit model, as a function of the illumination irradiance, with both full field and 1-pixel wide rectangular gratings for devices with (a) 100 µm bipolar pixels and (b) 40 µm monopolar pixels.

## IV. Conclusions and Impact

By using a sparse approximation to the original conductance matrix along with a low rank compensation matrix that scales linearly with the number of pixels, we unlock the full potential of RPSim for modelling neurostimulating arrays, including the next generation of subretinal implants. We see that in practice, with realistic stimulation patterns and parameters mimicking what is used in clinical settings (letters and sparse bars), the error resulting from this simplification is negligible - well below 1%, while reducing the netlist size and computation time by nearly 10-fold. Further reductions in computational length are also feasible for simulations involving larger implants. Speeding up the computation so dramatically will enable the use of iterative optimization techniques for the design of multi-electrode arrays operating under various conditions. This is important as electrode design plays a crucial role in the selectivity and efficacy of neurostimulation implants. Improper electrode design can result in poor stimulation of neurons, low contrast of the patterns, or the undesired stimulation of non-target cells. Designing electrodes for selective stimulation of cells is the key to achieving high resolution prosthetic vision, as well as other high-fidelity electro-neural interfaces.

## Acknowledgment

Authors declare no financial interest in the subject of this paper.

We would like to thank Yueming Zhuo and Ken Kundert for discussions on topics relevant to this paper.

